# Prediction of amyloid β accumulation from multiple biomarkers using a hierarchical Bayesian model

**DOI:** 10.1101/2022.10.21.513271

**Authors:** Yuichiro Yada, Honda Naoki

**Author notes:** Corresponding author: Honda Naoki, Address: Graduate School of Integrated Sciences for Life, 1-3-1 Kagamiyama, Higashi-Hiroshima City, Hiroshima, 739-8526, Japan. Co-corresponding author: Yuichiro Yada, Address: Graduate School of Integrated Sciences for Life, 1-3-1 Kagamiyama, Higashi-Hiroshima City, Hiroshima, 739-8526, Japan.

## Abstract

Accumulation of amyloid-beta (Aβ) in the brain is associated with neurodegeneration in Alzheimer’s disease and can be an indicator of early disease progression. Thus, the non-invasively and inexpensively observable features related to Aβ accumulation are promising biomarkers. However, in the experimental discovery of biomarkers in preclinical models, Aβ and biomarker candidates are usually not observed in identical sample populations. This study established a hierarchical Bayesian model that predicts Aβ accumulation level solely from biomarker candidates by integrating incomplete information. The model was applied to 5×FAD mouse behavioral experimental data. The predicted Aβ accumulation level obeyed the observed amount of Aβ when multiple features were used for learning and prediction. Based on the evaluation of predictability, the results suggest that the proposed model can contribute to discovering novel biomarkers, that is, multivariate biomarkers relevant to the accumulation state of abnormal proteins.

## Introduction

Alzheimer’s disease (AD) is the most common cause of dementia^1,2^. AD is a neurodegenerative disease in which neurons in the brain gradually die, causing progressive cognitive decline characterized by memory loss, impaired judgment and reasoning skills, communication difficulties, and changes in personality and behavior. However, most cases of AD progress sporadically, and many patients have no family history of AD^3^. Thus, the genetic background of sporadic AD has been extensively investigated, revealing the complex genetic architecture of late-onset neurodegenerative diseases^4–7^.

The diagnosis or risk foresight of AD before initiating the decisive progression of neuronal loss may enable the potential treatment of the disease or the administration of appropriate symptomatic medication. The rapidly growing population of affected people raises the urgent demand for developing novel conveniently observed biomarkers and prediction methods for AD before this time point^8–10^. In AD, the gradual accumulation of abnormal proteins, amyloid beta (Aβ), and tau precede irreversible neurodegeneration^1,2,11^. Aβ accumulates earlier, followed by tau. Abnormal proteins may propagate from the brain to other regions^12–14^. Thus, the accumulation level of abnormal protein forms may represent the progression stage of neurodegenerative diseases in the early phase^15–17^. The imaging of Aβ in the brain using positron emission tomography (PET) scan and its assessment in cerebrospinal fluid (CSF) potentially facilitate AD onset prediction^1,2,18–20^. Yet, the demand for more inexpensive and non-invasive methods to predict the risk of onset has led to further investigation of novel biomarker candidates in humans and preclinical models^21–23^.

To discover new AD biomarkers using animal models, researchers usually conduct statistical testing to detect differing features between the AD model and healthy wildtype samples at the same age. Progression timing and speed of pathology, however, may be heterogeneous even among AD model animals with the same genetic background. Such heterogeneity could mask the differences researchers aimed to detect (**Fig. 1A**). The accumulation level of Aβ in the brains of individual animals, which may reflect the progression state of AD pathology, may have been overlooked in past biomarker discovery in model animals. Ideally, time-series paired data of abnormal protein levels and multidimensional biomarkers are required to identify novel biomarkers regarding the relevance of Aβ accumulation in AD (**Fig. 1B**). However, longitudinal measurements of individuals are impossible due to animal sacrifices; data are obtained as snapshots (**Fig. 1C**). Another problem is that most studies have only measured either aberrant proteins or multidimensional biomarkers; few studies have measured both from enough samples (**Fig. 1C**).

**Fig. 1:**
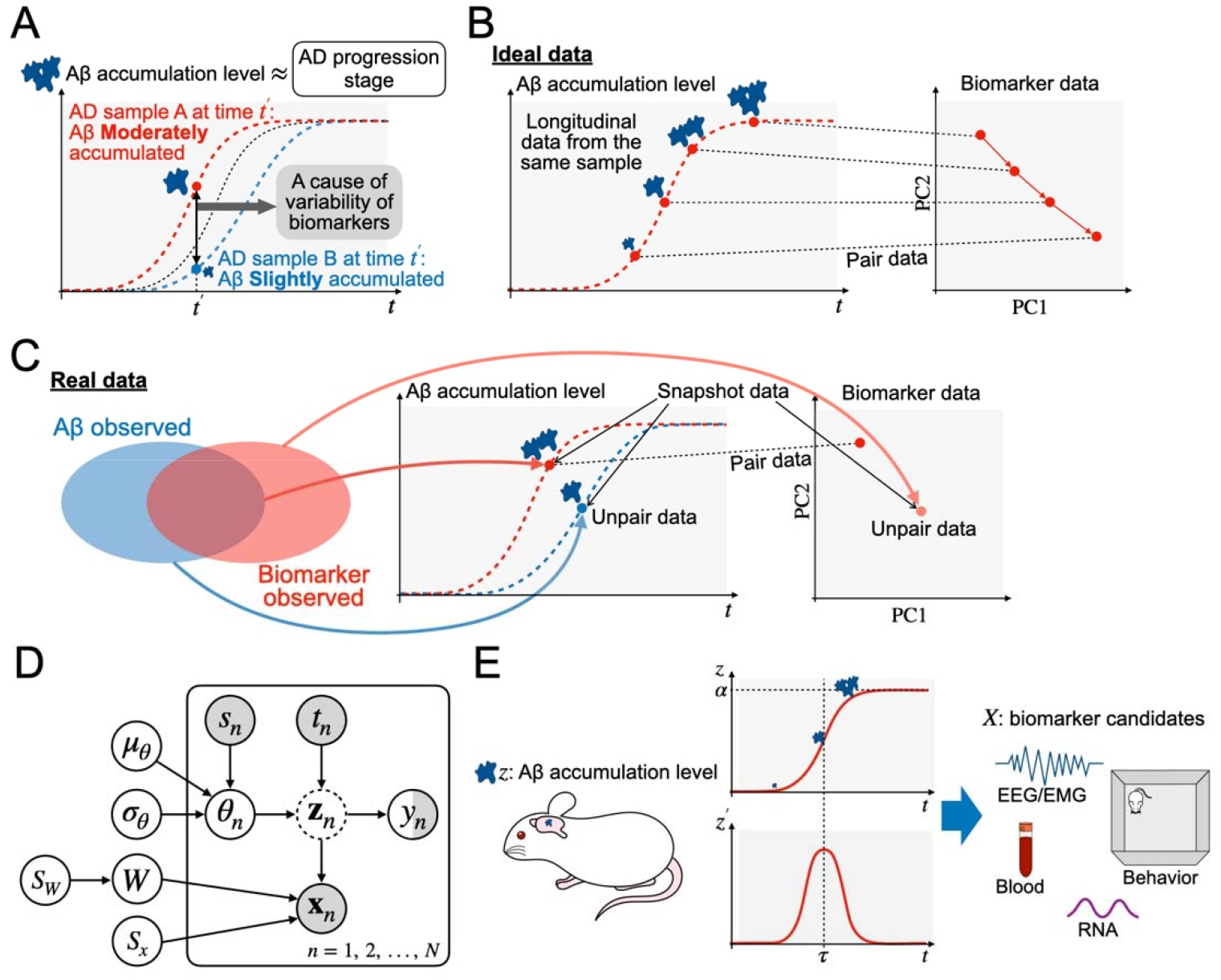
Concept and graphical model of the proposed model. (A) Alzheimer’s disease (AD) model samples (A and B) at the same age may have unignorably different amounts of accumulated amyloid beta (Aβ), which may correlate with the progression stage of early AD. (B) Ideally, Aβ-accumulation-relevant biomarkers are estimated from the longitudinal paired data of Aβ observation and biomarker candidate observation simultaneously. (C) In a real experiment, Aβ and biomarker candidates are usually observed primarily in distinct groups of samples (unpaired data). The limited number of samples may have paired data. Each sample has data from a single time point (snapshot). (D) The graphical model representation of our proposed model. (E) The concept of our model. Aβ accumulates in a particular brain region according to a logistic function of time. Observed data of biomarker candidates are generated depending on the accumulation level and the instantaneous accumulation speed of Aβ in the brain.

In the present study, to find Aβ-accumulation-relevant biomarkers overcoming the problem of limited data, we proposed a hierarchical Bayesian model that describes the Aβ accumulation process and observation of biomarker candidates. In the model, an abnormal form of Aβ deposits over time according to the logistic function, whose parameters are unique to each sample. Owing to the Bayesian probabilistic formulation, the model can naturally integrate unpaired and partially incomplete data, increasing the accumulation level’s predictability from the observed biomarkers. By applying the model to the behavioral data sets of public 5×FAD mice^24^, we predicted the accumulation level of Aβ solely from behavioral data, supporting the concept of biomarker discovery based on predictability.

## Results

### The hierarchical Bayesian model of Aβ accumulation

To represent the pathogenesis of AD in animal models, we developed a mathematical model describing the accumulation process of Aβ in the brain and the observation process of Aβ and biomarker candidates. In the model (**Fig. 1D**), Aβ accumulates over time according to the logistic function:

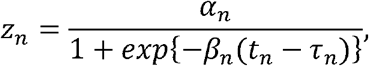

where *θ_n_* = {*α_n_*, *β_n_*, *τ_n_*} is a set of parameters depending on the individual animals; *α_n_*, *β_n_*, and *τ_n_* denote the maximum level of Aβ accumulation, steepness of Aβ accumulation, and critical period to reach half of the maximum Aβ, respectively. To express heterogeneity in the Aβ accumulation process among animals, different values of *θ_n_* = [*α_n_*, *β_n_*, *τ_n_*] are assigned to the individual animals, following the distribution shared within the same type of animals *s_n_*, that is, wildtype or AD model animals.

The observed amount of Aβ, *y_n_*, is subject to noise as *y_n_* = *z_n_* + *σ_y_ξ*, where *σ_y_* and *ξ* indicate the noise strength and Gaussian noise with zero mean and unit variance, respectively. The observed data of biomarker candidates 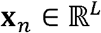 were assumed to reflect *z_n_* and its temporal derivative *z*′_*n*_, i.e., *dz_n_*/*dt*, as

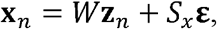

where **z**_*n*_ = (*z_n_*, *Cz*′_*n*_, 1)^T^, 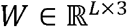 indicates weight matrix, *S_x_* = {*σ*_*x*_1__, *σ*_*x*_2__,′, *σ*_*x_L_*_) and 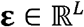 indicate Gaussian independent noises with zero mean and unit variance (**Fig. 1D**). *C* is a scaling factor common across samples. Here, we assumed that biomarker candidate data **x**_*n*_ was generated (**Fig. 1E**) via the same biological process among animals. *W* is thus the shared parameter among the animals sampled from the same distribution. This model was formulated based on a hierarchical Bayesian model (see Methods).

### Semi-supervised procedure to predict Aβ accumulation

Based on this model, we aimed to estimate the latent Aβ accumulation *z_n_* from the observed data of the biomarker candidate **x**_*n*_. To this end, we also needed to train the model by estimating the parameters from the data. Here, we made three assumptions, following the actual situation of available data: first, animals are sampled once as snapshots, not as time series, due to the requirement of euthanization. Second, most of the samples only had either Aβ accumulation or biomarker candidates, meaning that these were observed in different animal populations. Third, only a subset of the samples had paired data for the biomarker candidate and Aβ accumulation at the same age. Therefore, there were three kinds of data: unpaired data of Aβ accumulation, unpaired data of biomarker candidates, and paired data containing both (**Fig. 1C**).

Given the above assumption, we proposed a Bayesian probabilistic treatment to train the model by integrating the paired and incomplete unpaired data in a semi-supervised manner (see Methods). In the first step, using only the unpaired dataset of Aβ accumulation, we pre-trained the model by estimating the distribution of the logistic function parameter *θ* for each of the wildtype and AD model animals (**Fig. 2**). Next, using the estimated distributions in the first step as prior knowledge, we inferred the distribution of all model parameters from the remaining datasets; the unpaired dataset of biomarker candidates was used for unsupervised learning, whereas the paired dataset was used for supervised learning.

**Fig. 2:**
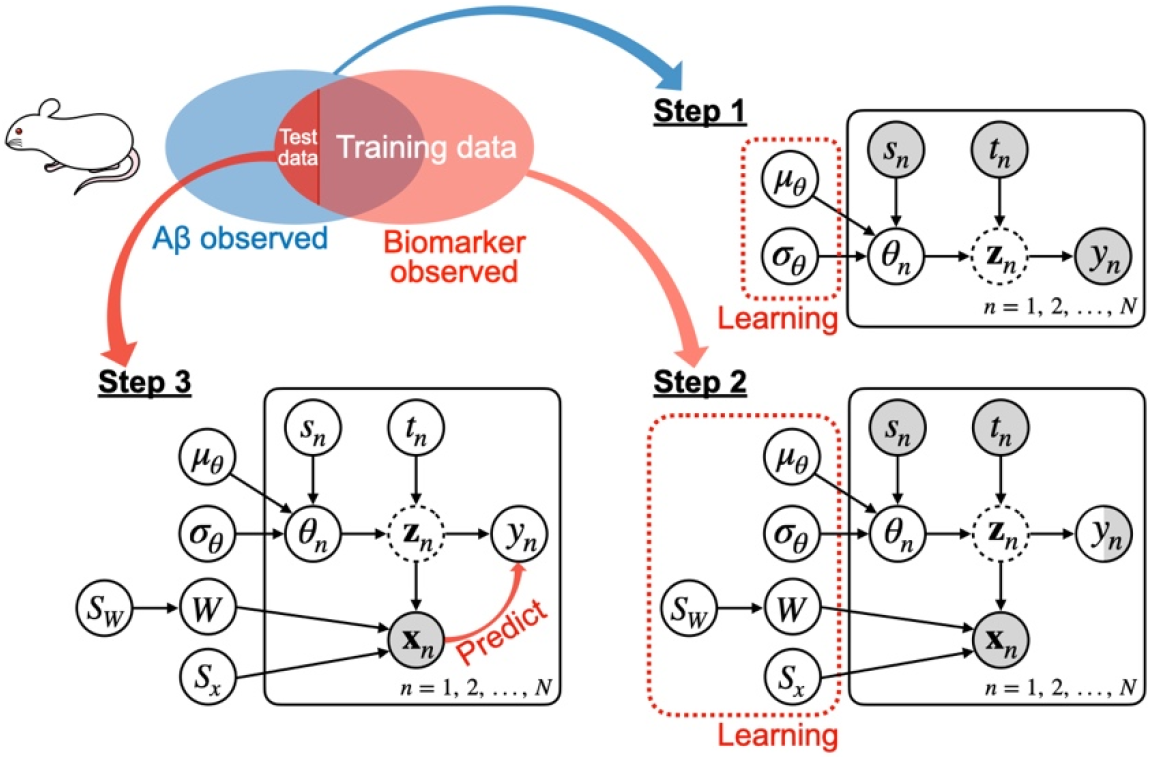
Learning and prediction steps of the proposed model. The model learns the parameters that integrate the incomplete data and predicts the Aβ accumulation from the observed data. In the first step, the distributions of the hyperparameters are inferred from the observed Aβ value. In the second step, the distributions of all parameters and hyperparameters were updated using the observed biomarker candidate data. Third, the predictability of the accumulated level of Aβ from the observed biomarker candidates was evaluated.

After learning the parameters, the model could predict the accumulated level of Aβ *z_n_* from the observed data of biomarker candidate *x_n_* (**Fig. 2**) (see Methods).

### Experiment data of AD model mice

As the application of our estimation method, we adopted real-world experimental data from the AD model and wildtype (WT) mice^24^. The AD model mice were 5xFAD transgenic mice with five human familial AD mutations in APP and PSEN1 on the background of a C57BL/6J strain, showing robust Aβ pathological accumulation and neuronal cell death. For AD model mice, most of the data contained unpaired data either of the Aβ accumulation (24 samples) or behavioral features (82 samples). In contrast, some paired data existed (18 samples only for 8- and 12-month-old mice) (**Fig. 3A**). Aβ accumulation was evaluated by the insoluble fraction level of Aβ40 and Aβ42 in the hippocampus of 42 AD model mice at 4, 8, 12, and 18 months of age (**Fig. 3D**). For WT mice, the observation of Aβ is unavailable in the dataset. We assumed that the insoluble Aβ in WT mice remained undetectable during the mouse’s lifetime based on the reports that WT mice do not develop Aβ plaque during their normal life span^25^. Each AD model and WT mouse were behaviorally evaluated; we selected three kinds of behavioral experiments where 11 features were obtained at 4, 8, or 12 months of age (Supplementary Table S1) (**Fig. 3B**). These 11-dimensional data were addressed as biomarker candidate data, and visualized using principal component analysis (PCA) (**Fig. 3C**).

**Fig. 3:**
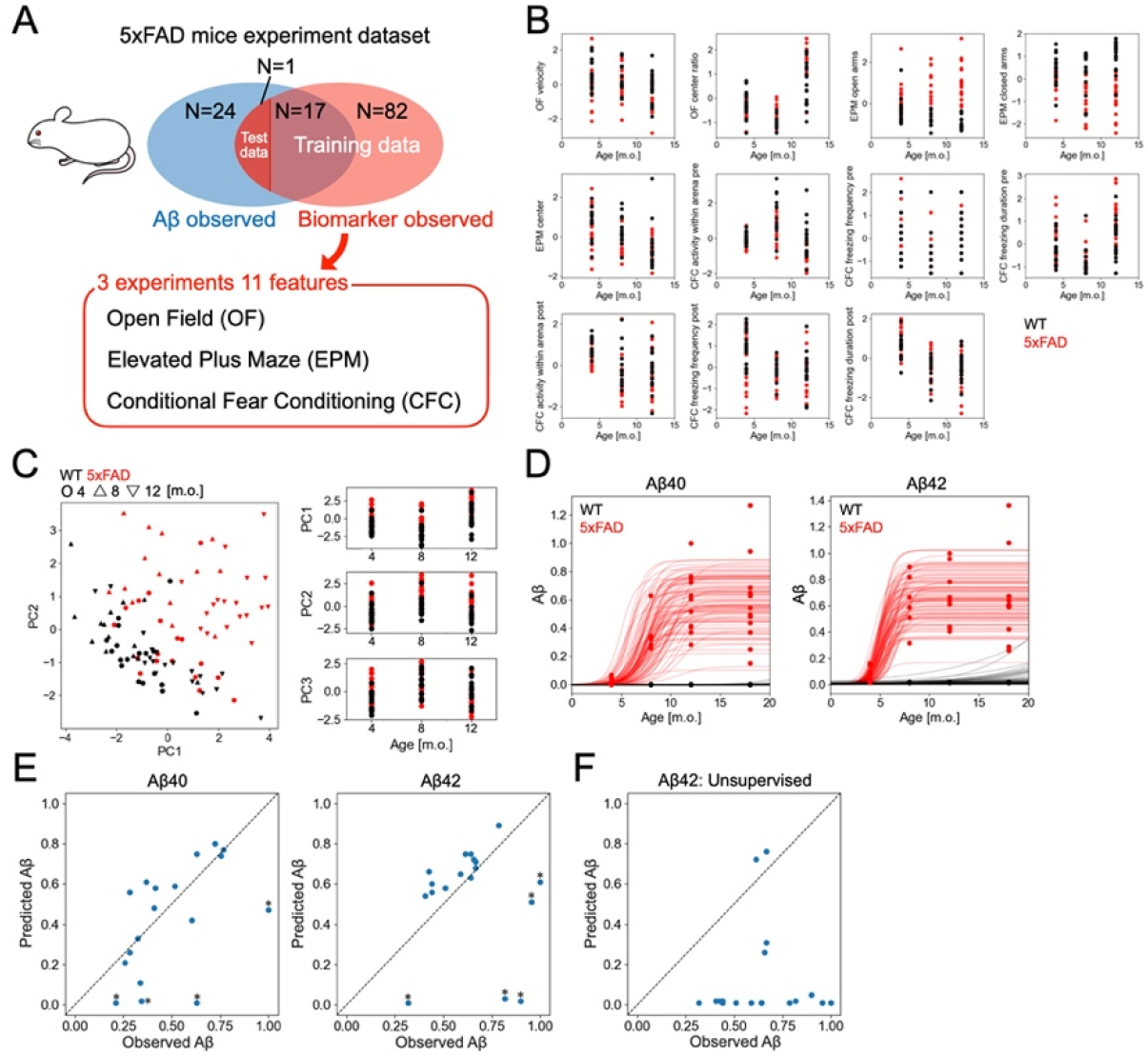
Application of the proposed model to 5×FAD mice data. (A) Components of the experimental data. The accumulation levels of Aβ were observed in 42 samples. The biomarker candidates, 11 features from 3 different behavioral experiments, were observed in 100 samples. The two groups partially overlapped: 18 samples have Aβ and biomarker candidate data. The predictability is evaluated by the leave-one-out cross-validation. (B) Time course of the behavioral features. The black points are from WT mice, and the red points are from 5×FAD mice. (C) PCA of the behavioral features obtained from the experiment. The red points correspond to 5×FAD samples, and the black points correspond to WT samples. In the scatter plot (left), the points’ shape represents the samples’ months of age (circle: 4, triangle: 8, inverted triangle: 12). (D) Inference of hyper-parameters of a logistic function. The data points are scaled insoluble fractions of Aβ_40_ and Aβ_42_ at the hippocampus. The red points are of 5×FAD mice, and the black points are of WT mice. The lines represent the example logistic functions with hyper-parameters randomly sampled from the learned distribution (red: 5×FAD, black: WT). (E) Accumulation levels of Aβ_40_ and Aβ_42_ at the hippocampus predicted by the trained model from the behavioral features. (F) The accumulation level of Aβ42 at the hippocampus predicted by the unsupervised-trained model.

### Prediction of Aß accumulation in AD model mice

Using our method, we predicted Aβ accumulation in the hippocampus using behavioral features as biomarker candidates. First, we pre-trained the model to learn the distributions of the parameters of the logistic function from the insoluble fraction level of Aβ. The pre-trained model generated logistic time courses, representing the observed insoluble fraction level of Aβ40 and Aβ42 (**Fig. 3D**). Next, we trained the model using an unpaired dataset of behavioral features and a paired dataset.

Using the trained model, we predicted the accumulation level of Aβ from the behavioral features of the paired data using leave-one-out cross-validation (**Fig. 3E**). The prediction errors were almost the same for the two types of Aβ (mean squared error (MSE)= 0.060 for Aβ40 and MSE = 0.111 for Aβ42). The majority of the predictions followed the observed amount of Aβ both in Aβ40 and Aβ42. However, several data were unpredictable with large errors (asterisks in **Fig. 3E**). Most of the unpredictable data were from 8-month-old mice and shared between Aβ40 and Aβ42. Moreover, 8-month-old mice showed larger prediction errors than 12-month-old mice (MSE of Aβ40 = 0.083 (8 months), 0.046 (12 months); MSE of Aβ42 = 0.214 (8 months), 0.046 (12 months)). If the training was implemented unsupervised, the prediction failed (**Fig. 3F**), indicating the importance of supervised information for prediction. The predictive performance for Aβ accumulation in the cortex was not as good as that in the hippocampus (Supplementary Fig. S1).

### Selection of biomarkers for prediction of Aß accumulation

To assess the importance of each behavioral feature in predicting Aβ accumulation, we removed each behavioral feature and evaluated the prediction error for each (**Fig. 4A**). We then found that predictive performance was considerably decreased by removing “time in the center” in the open field experiment and “time spent in the open arm” in the elevated plus maze experiment, which exhibited significant differences in WT and 5×FAD mice^24^. Next, we ranked the features by their impact on the prediction error and assessed the prediction performance by including features individually from the top of the ranks (**Fig. 4B**). The prediction error decreased considerably until the top five features of the ranks were recruited, which were composed of features from three different experiments. The results show that multivariate features from different experiments could be potential AD biomarkers.

**Fig. 4:**
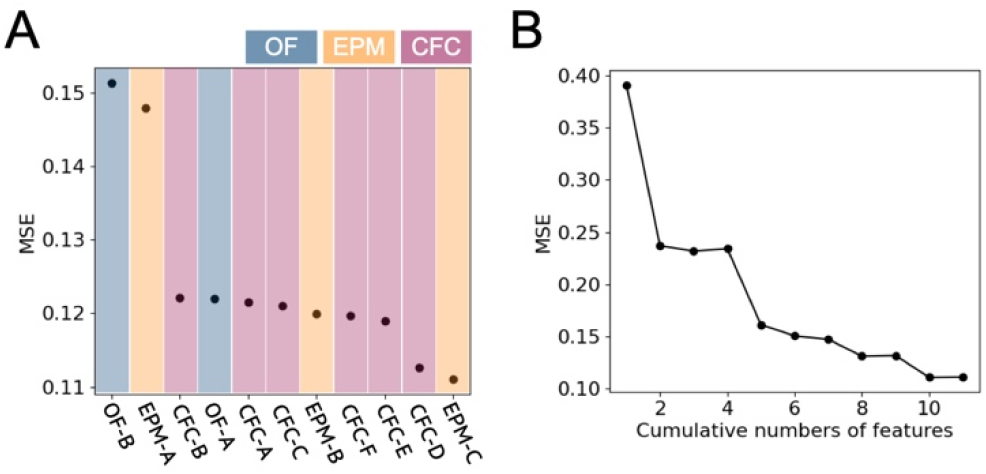
Evaluation of the behavioral features based on the predictability of the Aβ accumulation level. (A) Prediction error in the case of excluding a particular behavioral feature from the observation. (B) Variation of the prediction performance when the number of the observed features increases, where the features are sequentially included from the top of the list in Fig. 4A.

We also evaluated the number of features required for the proposed model to determine its predictability. Here, we randomly selected several behavioral features by varying their number, trained the model, and made a prediction (**Fig. 5A**). Statistically significant differences were detected between the predictions using 1–7 and 11 features (Mann–Whitney U-test, p<0.05). In contrast, there was no significant difference between the predictions using 10 and 11 features (Fig. 5B), suggesting that as many diverse features as possible are preferable to achieve better prediction performance.

**Fig. 5:**
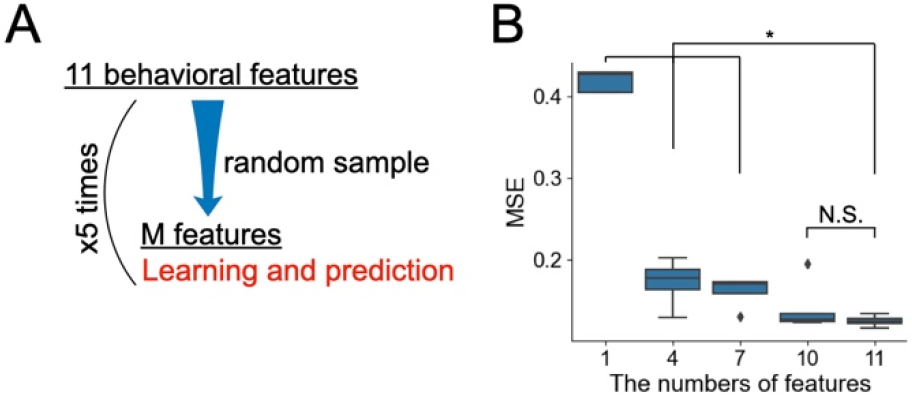
Learnings and predictions in randomly sampled features. (A) *M* features are randomly sampled from 11 behavioral features; the model learns the data using the *M* features, and the prediction performance is evaluated. (B) Mean squared errors between the observed amount of Aβ (Aβ_42_ at the hippocampus) and the predicted amount using *M* = {1, 4, 7, 10, 11} features for the learning and prediction. *M* features were sampled five times. (*p<0.05. Mann–Whitney U-test).

### Prediction of Aß accumulation by usual machine learning methods

The predictive performance of the proposed model was compared with that of standard machine learning techniques, namely ordinary linear and random forest regressions. We then demonstrated that our proposed model outperformed standard machine learning techniques both in prediction of Aβ40 and Aβ42 in the hippocampus (**Fig. 6A–D**). Indeed, the proposed model showed a smaller median absolute prediction error than that of the standard models (**Table 1**).

**Fig. 6:**
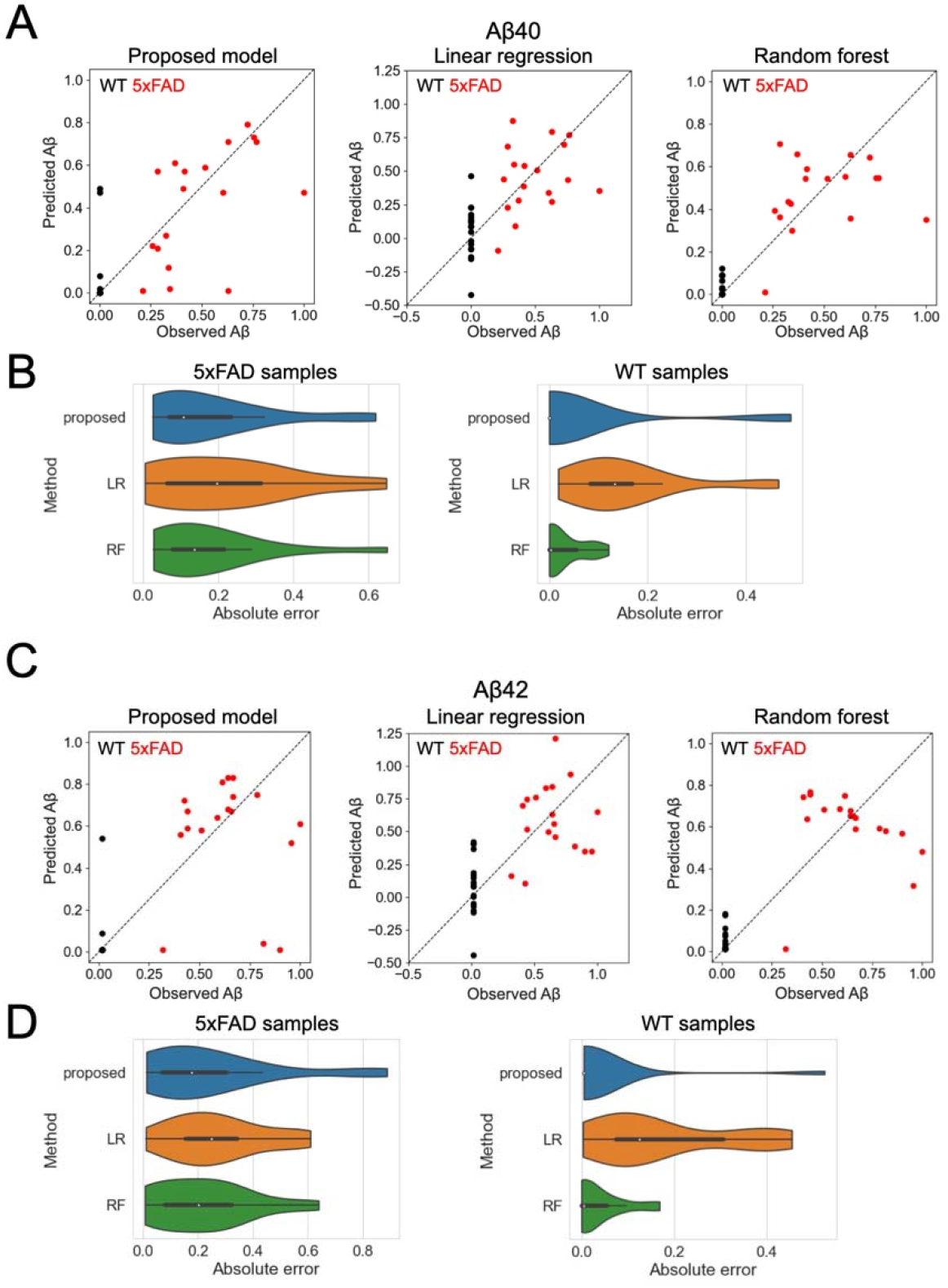
Comparison with standard machine learning methods. (A, C) Predictive performances against (A) Aβ40 and (C) Aβ42 of the proposed model, linear regression, and random forest regression. Eighteen virtually prepared supervised WT samples were used as test samples; the Aβ accumulation levels of such samples were assumed to be minimum quantity in the observation of Aβ. Three 5×FAD mice and three WT mice were used as test samples and evaluated with 6-fold cross-validation. (B, D) Distributions of absolute prediction errors when the accumulation levels of (B) Aβ40 and (D) Aβ42 were predicted for 5xFAD samples (left) and WT samples (right).

**Table 1:** Comparison of predictive performance with standard machine learning techniques.

| Method | Aβ | Sample type | Median absolute prediction error |
| --- | --- | --- | --- |
| Proposed | A$\beta_{40}$ | 5xFAD | 0.108 |
|  |  | WT | 0.0 |
| | A$\beta_{42}$ | 5xFAD | 0.177 |
|  |  | WT | 0.005 |
| Linear regression (LR) | A$\beta_{40}$ | 5xFAD | 0.197 |
|  |  | WT | 0.133 |
| | A$\beta_{42}$ | 5xFAD | 0.249 |
|  |  | WT | 0.125 |
| Random forest (RF) | A$\beta_{40}$ | 5xFAD | 0.136 |
|  |  | WT | 0.003 |
| | A$\beta_{42}$ | 5xFAD | 0.208 |
|  |  | WT | 0.005 |

### Application to synthetic data

In the 5×FAD experimental data used in this study, the paired data with Aβ accumulation and behavioral features were limited only to the phase when the accumulation process of Aβ has vastly progressed, that is, 8- and 12-month-old mice. Thus, the predictive performance of the proposed model for the earlier phase of Aβ accumulation remains unknown. To evaluate this, the model was applied to synthetic data that contained earlier phase samples.

To this end, we prepared synthetic unpaired and paired data at various ages, including the early phases, by simulating the model. The synthetic data were composed of 20 samples of Aβ accumulation alone, 50 samples of biomarker candidates alone, and 50 samples paired for the AD model and WT (**Fig. 7A**). First, we pretrained the model using the synthesized unpaired data of Aβ accumulation, representing the variation in the observed accumulation level of Aβ (**Fig. 7B**). We then trained the model using unpaired data of biomarker candidates and paired data using semi-supervised learning. We confirmed that the estimated parameters followed the ground truth used for the synthesized data (**Fig. 7C**), indicating that our estimation method was efficient. Using the trained model, we predicted *z*\*, the Aβ accumulation level of unknown samples, from the observed biomarker features *X*\* (**Fig. 7D**, top right). We evaluated the prediction accuracy by changing the ratio of supervised paired samples for training. The MSE changed slightly with varying ratios, suggesting that not many supervised samples were required for prediction (right and bottom left in **Fig. 7D**).

**Fig. 7:**
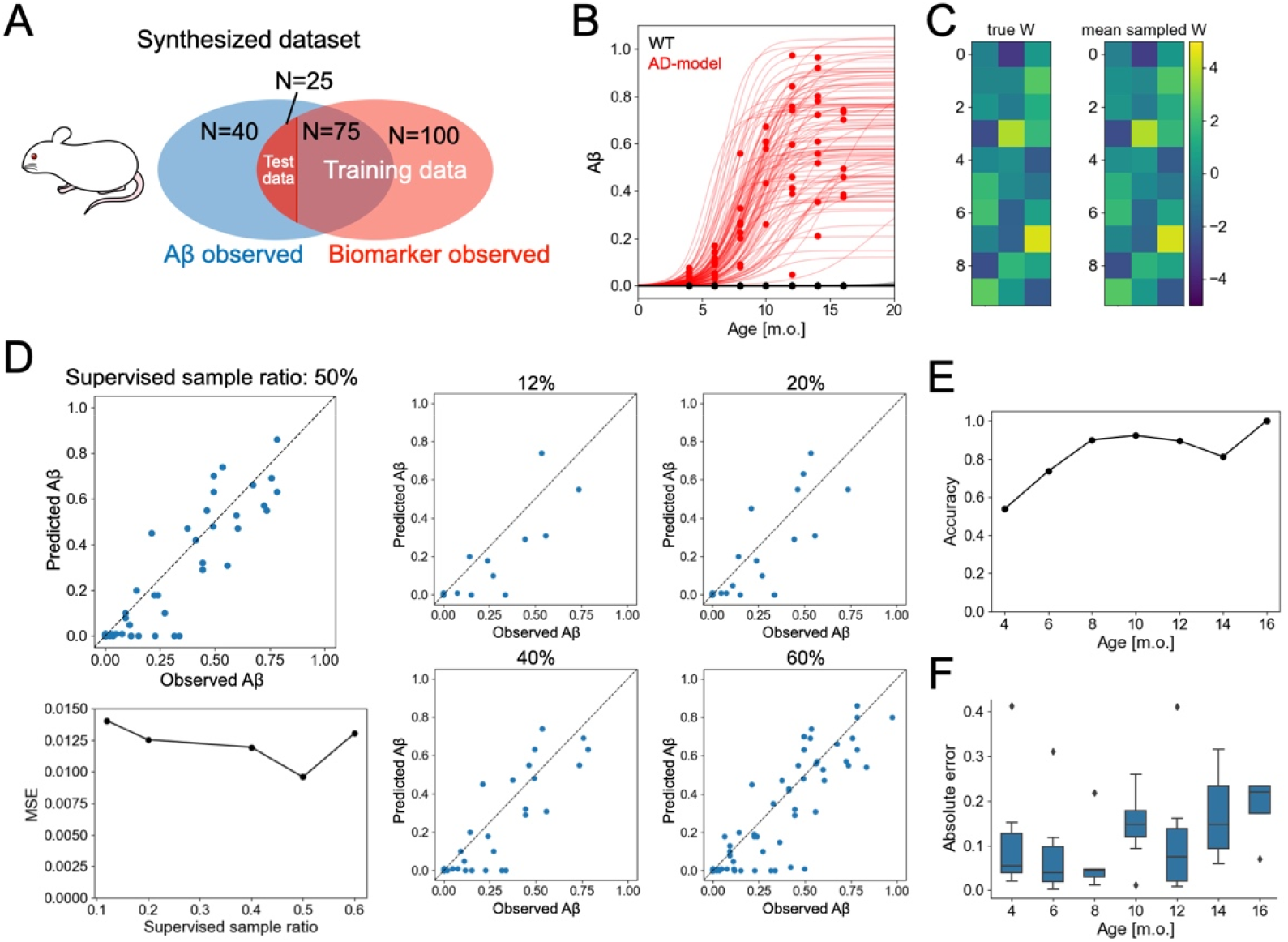
Learning and prediction performance of the proposed model is evaluated by synthesized data. (A) Components of the synthesized data. The accumulation levels of Aβ were observed in 140 samples. The biomarker candidates were observed in 200 samples. The two groups partially overlap: 100 samples have Aβ and biomarker candidate data. The ages of the samples are randomly sampled from *t_n_* = {4, 6, …, 16} and WT and AD model samples are in the same proportion. The predictability is evaluated by 4-fold cross-validation. (B) Inference of hyper-parameters of a logistic function from the observed amount of Aβ. The red points are the observed value of AD model samples, and the black points are those of WT samples. The example logistic functions with hyper-parameters are randomly sampled from the learned distribution are represented with red (AD model) and black (WT) lines. (C) Representative result of the inference of the model parameters from synthetically generated biomarker data. The left is true generated *W*, and the right is the mean of the posterior samples of *W*. (D) Prediction of the accumulation level of Aβ from the observed biomarker data by the learned model. The mean prediction errors according to the ratio of supervised samples (bottom left) and the prediction results at the respective conditions (right). (E, F) Accuracy of predicting the type of the samples (E) and absolute prediction errors in the AD model samples (F) according to the age of the samples.

Finally, we evaluated the predictive performance dependency of the samples on age. The discrimination of sample types, such as WT and AD model mice, were relatively accurate in the older samples compared to that in the younger ones (**Fig. 7E**). The predictive performance of Aβ accumulation in the AD model samples did not show a significant change according to age (**Fig. 7F**). The results suggest that the model can predict Aβ accumulation levels in samples in the early phase of the accumulation process, while the types of samples are sometimes mispredicted.

## Discussion

This study demonstrates the proof-of-concept of Aβ-predictability-based multivariate AD biomarker discovery. We proposed a hierarchical Bayesian model that describes how biomarker candidates are generated in response to the accumulation of Aβ in the hippocampus. Using this model, we predicted Aβ accumulation from the behavioral features obtained in the 5×FAD behavioral experiments. Based on the effect of each biomarker candidate on predictability, we revealed that multiple behavioral features from three different experiments could be important biomarkers for predicting Aβ accumulation level.

The proposed model can naturally integrate the information from paired and unpaired data. The inference of the distribution of the logistic function parameters from Aβ-observed unpaired data constrains the dynamic range of Aβ accumulation level. The information from unpaired data that lack an observed amount of Aβ provides the generation process of biomarker candidates and their variability. Learning information from the paired supervised data further helps to adjust the generation process. Notably, as the simultaneous observation of biomarker candidates and Aβ from the same sample is labor-intensive, expensive, and technically challenging, most samples usually lack information on either biomarker candidates or Aβ. In this situation, the proposed model makes it possible to make such incomplete datasets available for Aβ predictability-based biomarker discovery.

The predictive performance of the proposed model was found to be inferior in young mice than in older mice. The nature of the analyzed biomarker candidates and the proposed model limitations would have regulated the results. The principal component space of the 5×FAD mouse behavioral test data showed little difference between WT and AD modelo mice in the younger samples. The observed data of biomarker candidates may be generated as a non-linear transformation from Aβ accumulation, in contrast to the linearity assumed in the proposed model. We confirmed whether the model with nonlinear transformation yields better prediction performance by adopting a 3-layer neural network at the generation from z to *X*. However, the prediction performance only improved slightly (data not shown).

In humans, the accumulation of Aβ may initiate 10–20 years before the recognized cognitive decline^1,2^. However, in the model animals, behaviors and cognitive abilities were altered before the saturation of Aβ accumulation. For example, 5×FAD mice had a decline in memory function before 6 months of age, when Aβ was still in the accumulation process ^24,26,27^. These facts suggest that a probabilistic model sensitive to an earlier stage of Aβ accumulation is preferable to discover biomarkers in model animals. Furthermore, the gradient term of accumulated Aβ at the generation of biomarker data in the model might bestow such specificity to the proposed model when analyzing data that contain phase-specific biomarkers.

Whether Aβ acts upstream in the cascade leading to cell death, as in the amyloid hypothesis, is controversial^28,29^. However, Aβ indeed accumulates in the early stages of AD. Even if there is no causal relationship between the accumulation of Aβ and AD progression, Aβ may be a useful precursor for predicting AD. Notably, studies have reported alterations in the phenotypes of AD model mice that appear earlier than the onset of Aβ accumulation^30^. Such lesions may allow for an earlier definition of the latent state of AD progression.

Machine learning approaches for identifying the latent progression states of AD have recently attracted attention. Probabilistic models, such as mixed effect^31–35^ and hidden Markov models^36–38^ found the latent trajectories of the disease progression from longitudinal data of the clinical cohorts in unsupervised learning. Our approach assumed the accumulation level of Aβ as “the latent progression state” and estimated the state using semi-supervised learning. The molecular biological observation of Aβ is challenging in humans. However, the proposed framework is potentially beneficial for discovering non-invasive convenient biomarkers^8,9,39,40^ that are relevant to the amount of PET-detected Aβ or CSF Aβ in human data.

In other neurodegenerative diseases, such as Parkinson’s disease, Lewy body dementia, multiple system atrophy, Huntington’s disease, amyloid lateral sclerosis, and frontotemporal lobar degeneration, abnormal proteins accumulate in specific brain regions, leading to neuronal death^41–43^. The modeling approach presented in this paper is also potentially applicable to such neurodegenerative diseases. The risk of neurodegenerative diseases is a growing concern in an aging society. Predictability-based biomarker discovery by the proposed model may contribute to identifying biomarkers that make available predictions and potential interventions for diseases.

## Methods

### The generative model of Aβ and biomarker candidates

In the model, Aß is assumed monotonically accumulated as the logistic function:

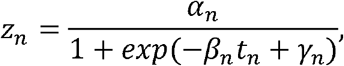

where *θ_n_* = {*α_n_*, *β_n_*, *γ_n_*} is a set of parameters, and *n* is the animal index. This equation can be rewritten from Equation (1), where *γ_n_* = *β_n_τ_n_*. Accordingly, the temporal derivative of the Aβ accumulation is

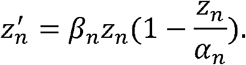

The parameters *θ_n_* depend on the individual, following distributions as

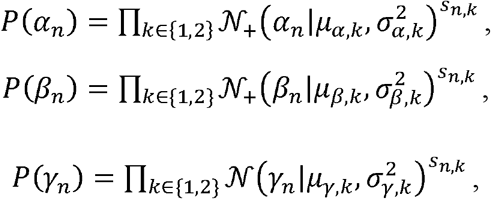

where 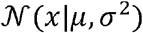 and 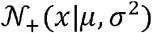 indicate normal distribution with mean *μ* and variance *σ*^2^ and truncated normal distribution with range of *x* > 0, respectively; **s**_*n*_ is one-hot vector representing the type of the samples, i.e., wildtype as (1, 0)^T^ or AD model mouse as (0, 1)^T^; *μ_ϕ,k_* and *σ*^2^_*ϕ,k*_ (*ϕ* ∈ {*α, β, γ*}) indicates parameters of wildtype (*k* = 1) or AD model mouse (*k* = 2).

The observed amount of Aß, *y_n_*, is generated from *z_n_* as

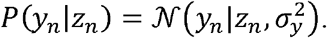

The observed data of *L*-dimensional biomarker candidates 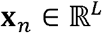 was generated from *z_n_* and 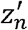 as

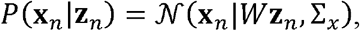

where **z**_*n*_ = (*z_n_*, *Cz*′*_n_*, 1)^T^, 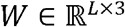 indicates weight matrix, and 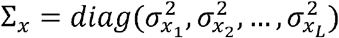 indicates a scaling factor common among samples that adjusts the range of *z*′.

### The prior distribution of parameters

For parameter estimation in a Bayesian manner, we introduced the prior distributions of parameters. The hyper-parameters of *P*(*α_n_*), *P*(*β_n_*) and *P*(*γ_n_*) are sampled from the following distributions:

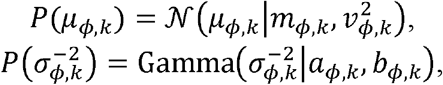

where *ϕ* ∈ {*α, β, τ*}, Gamma(*x*|*a, b*) indicates a Gamma distribution with shape parameter *a* and rate parameter *b*.

The prior distributions of *W* was

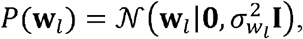

where **w**_*l*_ indicates the *l*-row of the weight matrix *W*. Prior of its hyper-parameter 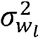 was hierarchically introduced as

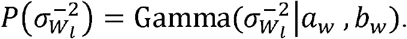

The prior distributions of 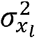 was

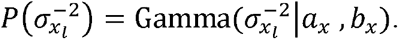

In the MCMC, we used 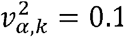, *a_α,k_* = 25, *b_α,k_* = 0.5, 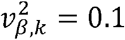, *α_β,k_* = 100, *b_β,k_* = 1, 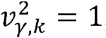, *a_γ,k_* = 10, *b_γ,k_* = 1 *v_α,k_* = 0.1, *a_αk_* = 25, *b_α,k_* = 0.5, *a_w_* = 100 *b_w_* = 1000 *a_x_* = 0.5 and *b_x_* 1 and 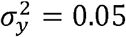.

When the model learned the distributions of the hyper-parameters from the Aß observation (step 1), we set *m_α,k_*, *m_β,k_* and *m_γ,k_* as the values estimated by the least square method.

### Bayesian inference of parameters

The model parameters were learned in two steps. In the first step (step 1), we inferred the posterior distributions of the parameters of a logistic function and those of hyper-parameters, given the data of Aβ accumulation as

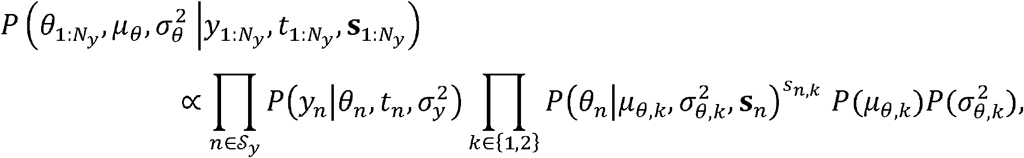

where *μ_θ_* {*μ*_*θ*, 1_, *μ*_*θ*, 2_}, *μ*_*θ, k*_ = {*μ_α, k_*, *μ_β, k_*, *μ_γ, k_*}, 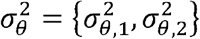, 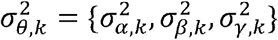, 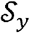 is the set of samples with Aβ accumulation in the training data, and *N_y_* is the number of samples in 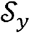. We assumed that the accumulation level of Aβ *z_n_* = 0 at time *t_n_* = 0 in all samples. This posterior distribution was estimated using the MCMC sampling algorithms. No-U-Turn samplers (NUTS) were used in this step because it is difficult to derive the closed form of the posterior distribution. From the posterior samples, we estimated the parameters of the Gaussian and Gamma distributions for *μ_θ,k_* and 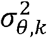, respectively, and these distributions with the estimated parameters were used as priors for the hyper-parameters. In the first step, MCMC sampling was performed across three independent chains, where 3,000 samples were drawn for burn-in, and another 3,000 were drawn to estimate the distribution.

In the second step (step 2), we inferred the posterior distributions of all parameters in the model, given the unpaired data of biomarker candidates and the paired data of Aβ accumulation and biomarker candidates. We used the NUTS-within-Gibbs approach for the inference. The weight matrix *W*, the variance of the weight matrix 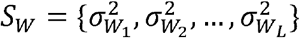, the variance of the observation noise *σ_x_* were sampled using Gibbs sampling as follows:

The Gibbs sampler for weight matrix *W*:

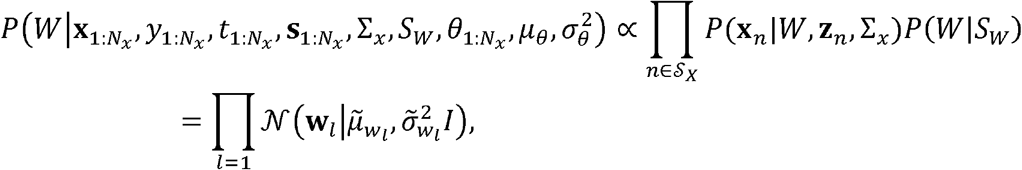

where 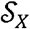 is a set of samples at least having biomarker candidates in the training data, *N_X_* is the number of samples in 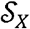, and

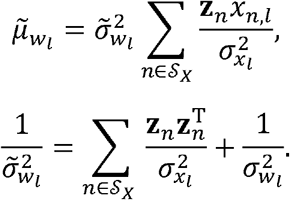

The Gibbs sampler for observation noise of biomarker candidates *S_x_*:

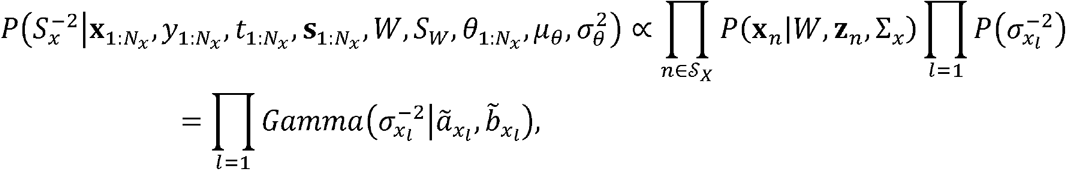

where

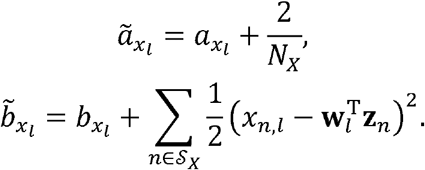

The Gibbs sampler for variance of the coefficient matrix *S_W_*:

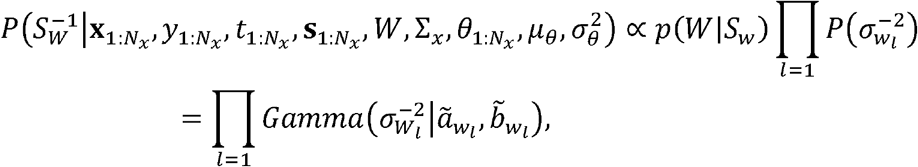

where

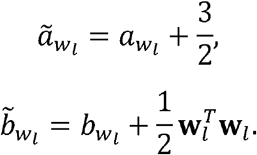

The posterior distribution of other parameters was sampled by MCMC using NUTS as

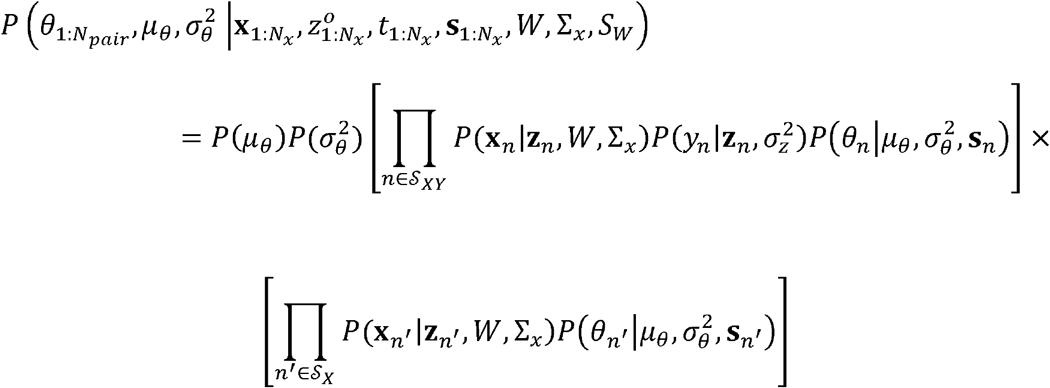

where 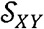 and 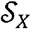 are sets of samples with paired data of Aβ accumulation and biomarker candidates and the unpaired data of biomarker candidates, respectively. We adopt the distributions estimated in Step 1 as *P*(*μ_θ_*) and 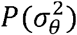. In the second step, MCMC sampling was performed across three independent chains, where 5,000 samples were drawn for burn-in, and another 5,000 were drawn to estimate the distribution. The inference program was implemented with Python using the NumPyro framework.

### Prediction of Aß accumulation from biomarkers

In the prediction of Aß accumulation in the test data (step 3), we computed a conditional posterior predictive distribution of *y*\* given **x**\* as the following equation:

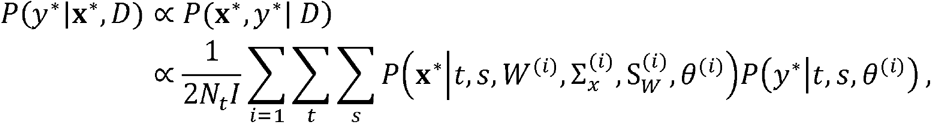

where *D* is the learned training data, *W*^(*i*)^, 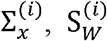, and *θ*^(*i*)^ are the posterior samples of the parameters, *I* is the number of the posterior samples and *N_t_* is the number of time points considered in the prediction *t* = {2, 3,…, 18}.

### Behavioral experiments with 5×FAD mice

We used the dataset from Forner et al.^24^, freely available in the public repository. Eleven features from three experiments were analyzed using our proposed model. In the open-field experiment, the velocity and time ratio in the center (the time in the center divided by the time in the arena) was used in the analysis. In the elevated plus maze experiment, the amount of time a mouse spent cumulatively in the open arm, closed arm, and center area of the maze was used in the analysis. In the contextual fear conditioning experiment, the activity level, inactive freezing frequency, and cumulative duration of inactive freezing were monitored for each mouse during a 2-min habituation and exploration in a chamber. Subsequently, electrical shock was applied to the mouse. After 24 h, the same behavioral features were monitored for 5 min in the chamber.

### Preprocessing

Preprocessing was performed to analyze 5xFAD mouse behavioral data. Behavioral features were standardized such that the mean of each feature was zero and the standard deviation of each feature was 1.0. The observed amount of Aβ was scaled so that the maximum observed value from 12 months of age mice equals 1.0. Based on the assumption that insoluble Aβ in the brain of WT mice remains undetectable throughout their lives, we virtually generated WT samples at 8, 12, and 18 months of age, where the observed amount of Aβ at each time sample was 0.0. A 5×FAD mouse “individual ID = 572” was excluded from the paired-data samples because the measurement of insoluble Aβ in the sample may have failed.

## Acknowledgments

This work is supported in part by Moonshot R&D–MILLENNIA Program [grant number JPMJMS2024-9] by JST.

## Author Contributions

Y.Y. and H.N. conceived the project and developed the method. Y.Y. implemented the software and analyzed data. Y.Y. and H.N. wrote the manuscript.

## Competing interests

The authors declare no competing interests.

## Data availability

Datasets for the current study are available from Forner et al., *Scientific Data*, 2021 (https://doi.org/10.1038/s41597-021-01054-y).

## Notes

### Competing Interest Statement

The authors have declared no competing interest.

### Summary of Updates

There are no significant differences between this version and the previous version.

