## Supplemental Fig. S1 and Table S1 for "Prediction of amyloid β accumulation from multiple biomarkers using a hierarchical Bayesian model"

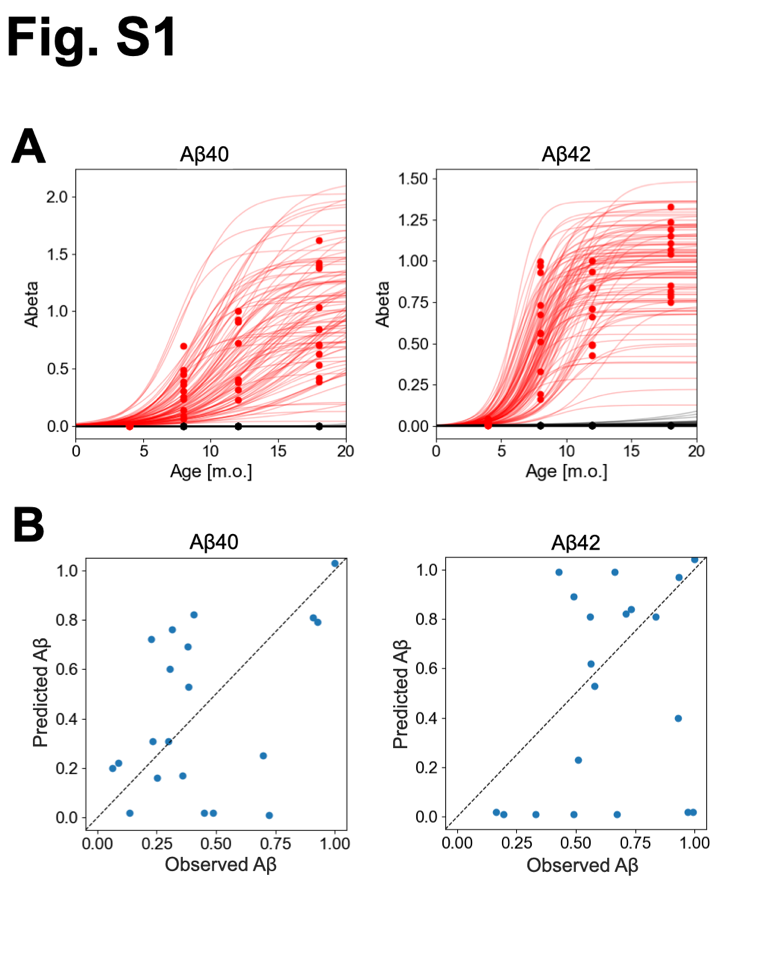

**Fig. S1: The results of the learning and prediction of Aß in the cortex.**

(A) The inference of hyper-parameters of a logistic function. The red points are for AD model samples and the black points are for WT samples. The lines represent the example logistic functions whose hyper-parameters are randomly sampled from the learned distribution (red: 5xFAD, black: WT). (B) The results in the predictions of accumulation levels of Aß_40_ and Aß_42_ at the cortex by the trained model.

Table S1: Behavioral experiments and features from Forner et al. used in this study.

| Behavioral experiment | Index | Feature |
| --- | --- | --- |
| Open field (OF) | A | Velocity |
|  | B | Time ratio in the center* |
| Elevated plus maze (EPM) | A | Cumulative open arms |
|  | B | Cumulative closed arms |
|  | C | Cumulative center |
| Conditional fear conditioning (CFC) | A | Baseline train activity within arena mean |
|  | B | Baseline train inactive freezing frequency |
|  | C | Baseline train inactive freezing cumulative duration |
|  | D | test activity within arena mean |
|  | E | test inactive freezing frequency |
|  | F | test inactive freezing cumulative duration |

*defined as “time in the center” / “time in the arena”.
